# Plakoglobin supports length-dependent force generation and cardiac resistance to fasting

**DOI:** 10.64898/2026.10.09.757844

**Authors:** Samrat Chakraborty, Nashwa Jubran, Stephanie M. Deditz, Jonathan A. Kirk, Shenhav Shemer

## Abstract

Mechanical activity protects muscle from atrophy, but how contractile forces relate to anabolic signaling remains unclear. We previously showed that plakoglobin promotes insulin-PI3K-AKT-FoxO signaling and that its loss causes skeletal muscle atrophy. Here, super-resolution microscopy revealed that plakoglobin localizes with desmin at the sarcolemma and is lost from the membrane during fasting-induced skeletal muscle atrophy. In skinned muscle fibers, plakoglobin downregulation impaired stretch-induced increase in myofilament activation at long sarcomere length. During fasting, skeletal muscle lost sarcolemmal plakoglobin-containing complexes, insulin signaling, and mass and induced MuRF1 and MAFbx/Atrogin1. In contrast, cardiac muscle retained these complexes and anabolic signaling, and maintained its mass. Cardiac plakoglobin also resisted AAV-shJUP-mediated depletion despite viral delivery. Moreover, MEK inhibition disrupted this cardiac response, reduced AKT signaling, induced atrogenes, and decreased cardiac mass. Thus, plakoglobin supports force generation at increased sarcomere length, and preservation of plakoglobin-containing complexes and MEK-ERK signaling accompany cardiac resistance to fasting-induced atrophy.

## INTRODUCTION

Skeletal muscle is the most abundant tissue in the human body, and the major site for glucose disposal and energy homeostasis. Preservation of muscle mass and function is therefore critical for health and survival (Zhou et al., 2010). Loss of muscle mass (i.e. atrophy) due to aging or disease reduces quality of life and contributes to mortality (Cohen et al., 2015). Mechanical activity is one of the most effective physiological mechanisms that protect muscle from most types of atrophy, yet the mechanisms that couple contractile forces to anabolic signaling remain unclear.

Plakoglobin (γ-catenin) was originally identified as a component of desmosomes and adherens junctions in tissues that must withstand mechanical stress, especially cardiomyocytes and epithelia (Cowin et al., 1986; Knudsen and Wheelock, 1992), In these tissues plakoglobin contributes to cell-cell adhesion and signaling pathways that regulate cell growth and differentiation (Yin et al., 2005; Barker and Clevers, 2000). More recently, plakoglobin has been identified as a component of skeletal muscle costameres (Eid Mutlak et al., 2020; Cohen et al., 2014; Murgia et al., 2022), membrane-associated structures that link the contractile apparatus to the extracellular matrix and transmit mechanical forces across the sarcolemma. We previously demonstrated that plakoglobin binds the insulin receptor and PI3K and promotes PI3K-AKT-FoxO signaling in skeletal muscle (Cohen et al., 2014). Plakoglobin also stabilizes the dystrophin glycoprotein complex (DGC)-insulin receptor assemblies on the muscle membrane, and its downregulation alone compromises these protein assemblies, reduces PI3K-AKT signaling and causes skeletal muscle atrophy (Eid Mutlak et al., 2020). In addition, plakoglobin is reduced in several forms of muscle wasting, including disuse (e.g., bed rest, microgravity during spaceflight), diabetes, fasting and Duchenne muscular dystrophy, leading to destabilization of DGC-insulin receptor assemblies and impaired anabolic signaling (Eid Mutlak et al., 2020; Murgia et al., 2022).

In cardiac muscle, plakoglobin is an essential component of intercalated discs where it mediates adhesion between cardiomyocytes and contributes to mechanical coupling of the contractile apparatus. Mutations in the plakoglobin gene (*JUP*) cause arrhythmogenic right ventricular cardiomyopathy (ARVC) (Li et al., 2011), a disease characterized by progressive loss of cardiac muscle and life-threatening arrhythmias (McKoy et al., 2000; Asimaki et al., 2009). These observations highlight the importance of plakoglobin for maintaining the integrity of mechanically active tissues. However, whether plakoglobin directly participates in mechanotransduction in muscle remains unknown.

Emerging evidence suggests that several signaling pathways involved in muscle growth are regulated by mechanical forces. The extracellular signal-regulated kinase (ERK) pathway can be activated by mechanical stretch, integrin signaling, and cytoskeletal tension, and contributes to the regulation of cell growth, motility and metabolism (Hirashima et al., 2023; Tanimura et al., 2016). In muscle, ERK signaling has been implicated in adaptive responses to mechanical load and contractile activity (Thomas et al., 2024; Hirata et al., 2015). These observations raise the possibility that mechanotransduction pathways cooperate with metabolic signaling to preserve muscle integrity during physiological stress (e.g. catabolic states, strenuous exertion).

Here, we show that plakoglobin supports force generation at increased sarcomere length and that the heart resists fasting-induced atrophy by preserving plakoglobin-containing membrane complexes and anabolic signaling. Skeletal muscle loses plakoglobin-containing membrane complexes, insulin signaling, and mass, whereas the heart preserves these complexes and remains relatively resistant to atrophy. We further show that cardiac plakoglobin resists AAV-shJUP-mediated depletion despite viral delivery, and that MEK-ERK signaling is required to maintain anabolic signaling and cardiac mass during fasting. These findings extend plakoglobin function beyond its established role in skeletal muscle insulin signaling and identify a mechanical response associated with preservation of cardiac muscle during catabolic stress.

## RESULTS

### Plakoglobin is reduced from the sarcolemma during fasting-induced skeletal muscle atrophy

Mechanical activity is essential for maintenance of muscle mass and function. Because plakoglobin is distributed along the skeletal muscle membrane, associates with the DGC and promotes insulin receptor signaling in muscle (Eid Mutlak et al., 2020; Cohen et al., 2014; Yatsenko et al., 2020; Murgia et al., 2022), we asked whether its localization changes under conditions that induce muscle wasting. To examine the subcellular distribution of plakoglobin in muscle fibers with high precision, we performed super resolution Structured Illumination Microscopy (SIM) on longitudinal mouse Tibialis Anterior (TA) muscle sections labeled with plakoglobin and desmin antibodies (Fig. 1A and B). In muscles from fed animals, plakoglobin appeared as punctate structures along the sarcolemma, as shown previously (Cohen et al., 2014; Eid Mutlak et al., 2020), and vertically within intracellular structures that colocalized with desmin (Fig. 1B and C). This pattern is consistent with localization at membrane-cytoskeleton interphases (Konieczny et al., 2008; Agnetti et al., 2022).

**Figure 1.**
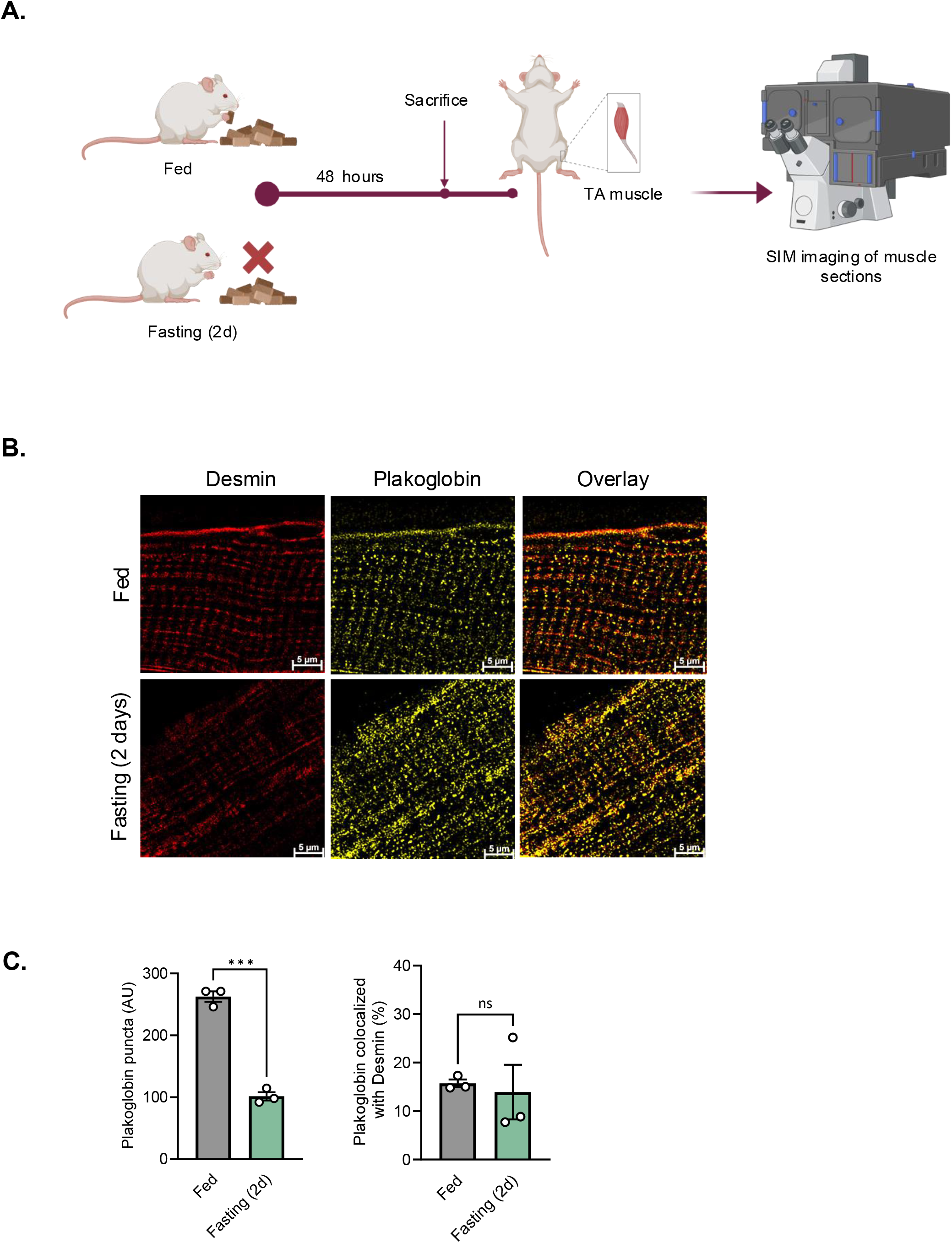
Muscle atrophy is associated with reduced plakoglobin membrane localization. (A) Schematic illustration of fasting induced atrophy and muscle analysis by SIM imaging. (B) Fasting reduces membrane-associated plakoglobin in skeletal muscle. Paraffin-embedded longitudinal sections of TA muscles from fed or fasted mice were stained with plakoglobin and desmin antibodies and imaged with SIM (bar, 5μm). Representative images are shown. n = 3 mice per condition. (C) Atrophy reduced sarcolemmal plakoglobin without altering its spatial association with desmin. Left: Average count of plakoglobin foci on the sarcolemma was quantified using the spots module of the Imaris software. n = 3 mice per condition, and 2 random areas were imaged per section (averaged values per biological replicate are shown). Data are mean ± SEM. ***, p<0.001 *vs.* fed by two-tailed unpaired Student *t* test. Right: Average colocalization of plakoglobin with desmin was quantified using Imaris software. Only spots within a distance threshold of < 100 nm were considered colocalized. n = 3 mice per condition, and 2-3 random areas were imaged per section (averaged values per biological replicate are shown). Data are mean ± SEM. ns, not statistically significant by two-tailed unpaired Student *t* test.

Because fasting is a well-established model of systemic catabolic state, which is also characterized by reduced mechanical loading due to weakness, we next examined plakoglobin localization after two days of food deprivation (Fig. 1A). Under these conditions, the number of plakoglobin puncta at the sarcolemma was markedly reduced (Fig. 1B and C). Similar reductions in plakoglobin membrane content have been observed in mouse models for type-2 diabetes, Duchenne muscular dystrophy, and in human muscle following disuse (bed rest) and microgravity (spaceflight) (Eid Mutlak et al., 2020; Murgia et al., 2022). However, the spatial association of the remaining plakoglobin with desmin was unchanged (Fig 1C, right), probably because desmin content also decreases in atrophying muscle (Aweida et al., 2018; Cohen et al., 2012; Vilchinskaya et al., 2015; Boudriau et al., 1996). Thus, plakoglobin associates with membrane structures in skeletal muscle fibers, and its loss from the sarcolemma during fasting may be a common feature of atrophy.

### Cardiac muscle preserves plakoglobin complexes and resists fasting-induced atrophy

We next asked whether the response of skeletal muscle to fasting differs in the heart, a muscle that contracts continuously. Fasting markedly reduced TA and Gastrocnemius (GA) muscle mass, whereas cardiac muscle mass remained similar to that in fed controls (Fig. 2A). Normalization to body weight (BW) produced an apparent increase in the cardiac muscle-to-body weight ratio because fasting reduced total body mass (Fig. 2B). This increase therefore does not indicate cardiac hypertrophy. In contrast, skeletal muscle mass remained significantly reduced even after normalization to BW (Fig. 2B). Thus, skeletal muscle undergoes substantial atrophy during fasting whereas cardiac muscle is relatively protected.

**Figure 2.**
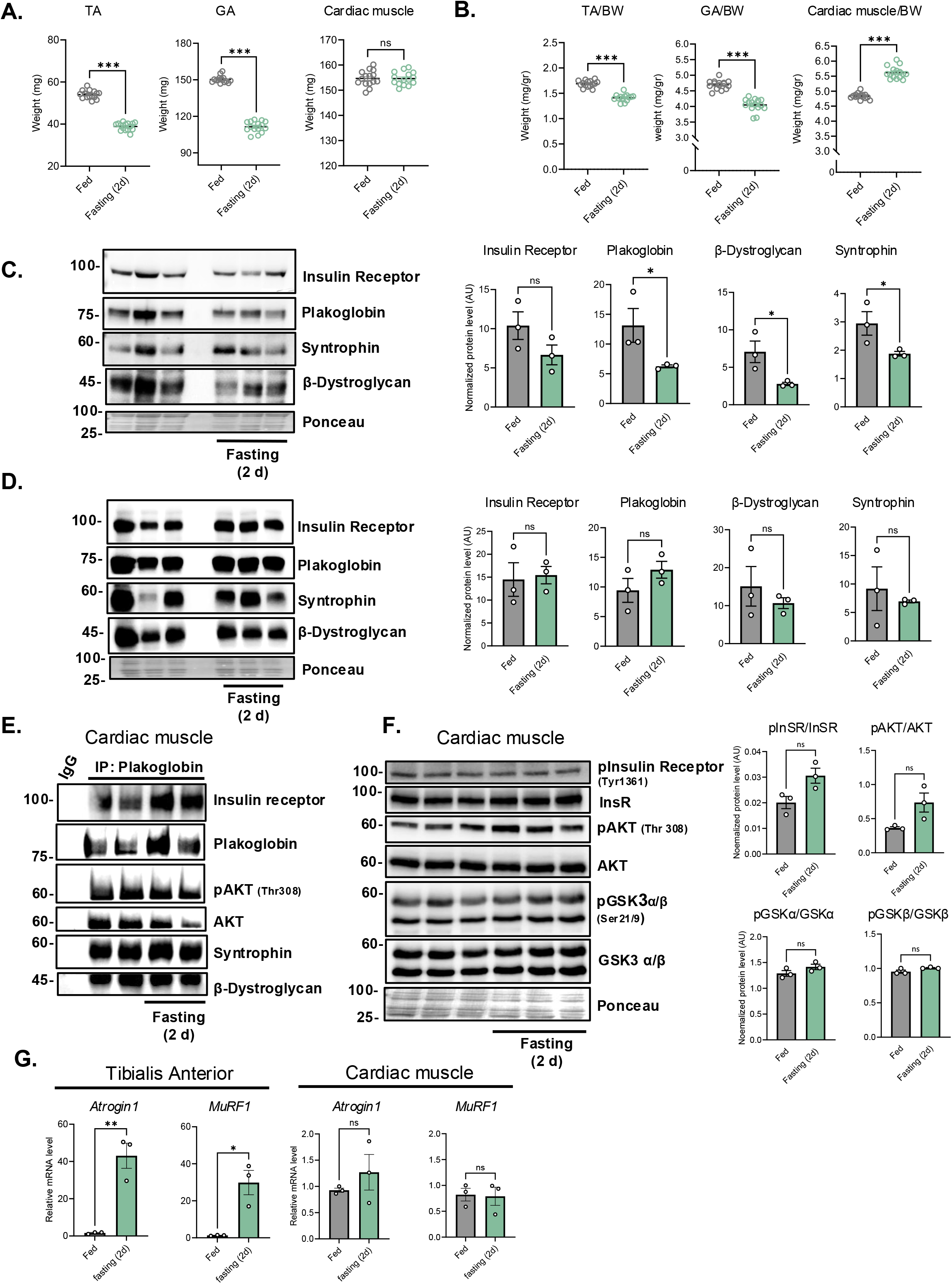
Cardiac muscle preserves plakoglobin complexes and resists fasting-induced atrophy. (A-B) TA, GA and cardiac muscle weights from fed and fasted (2 d) mice are shown either as absolute weights (mg) or normalized to body weight (mg/g BW). Mice had comparable body weights at the start of the experiments. Data from three independent experiments were pooled, yielding n =15 mice per condition. Data are mean ± SEM. ns, not statistically significant. ***, p<0.001 *vs*. fed by two-tailed unpaired Student *t* test. (C) During fasting, plakoglobin-containing protein assemblies are reduced in TA muscle membrane. Left: membrane extracts from TA muscles were analyzed by immunoblotting. Right: densitometry analysis of presented blots. n =3 mice per condition. Data are presented as mean ± SEM. ns, not statistically significant. *, p<0.05 *vs*. fed by one-tailed unpaired Student *t* test. (D) During fasting, plakoglobin-containing protein assemblies remain intact in cardiac muscle membrane. Left: membrane extracts from muscles were analyzed by immunoblotting. Right: densitometry analysis of presented blots. n =3 mice per condition. Data are presented as mean ± SEM. ns, not statistically significant *vs*. fed by two-tailed unpaired Student *t* test. (E) The heart maintains membrane-associated plakoglobin signaling complexes during fasting. Plakoglobin was immunoprecipitated from membrane extracts from cardiac muscles from fed or fasted mice, and protein precipitates were analyzed by immunoblotting. Mouse non-specific IgG was used as a control. n = 2 mice per condition. (F) Insulin signaling is preserved in cardiac muscle during fasting. Soluble fractions from cardiac muscles from fed or fasted mice were analyzed by immunoblotting. n = 3 mice per condition. Data are mean ± SEM. ns, not statistically significant *vs*. fed by two-tailed unpaired Student *t* test. (G) During fasting, atrophy-related genes are induced in skeletal muscle, but not in the heart. RT-PCR of mRNA preparations from cardiac or TA muscles from fed and fasted mice using specific primers for MAFbx/Atrogin1 or MuRF1. n = 3 mice per condition. Data are mean ± SEM. ns, not statistically significant. *, p<0.05 and **, p<0.01 *vs*. fed by two-tailed unpaired Student *t* test.

This difference was accompanied by preservation of plakoglobin signaling complexes in the heart. Immunoblot analysis of membrane extracts from mouse TA muscles revealed that fasting reduced the abundance of the DGC components β-dystroglycan and syntrophin, as well as plakoglobin, consistent with our prior observations (Eid Mutlak et al., 2020). The insulin receptor showed a trend of reduction although this trend did not reach significance (Fig. 2C). In contrast, we detected no fasting-induced reduction in the membrane content of these proteins in the heart (Fig. 2D). Co-immunoprecipitation of plakoglobin from membrane extracts of cardiac muscles confirmed that plakoglobin remained associated with the insulin receptor and DGC components during fasting, to a similar extent as in fed controls (Fig. 2E). In addition, the downstream insulin signaling effector AKT coprecipitated with this protein assembly in its active phosphorylated form (Thr308) (Fig. 2E). Consistently, insulin-AKT-GSK3 signaling remained active in cardiac muscle during fasting, despite the low circulating insulin levels, because the abundance and phosphorylation of insulin receptor and downstream signaling effectors, including phospho AKT (Thr 308) and phospho GSK3α/β (Ser21/9), did not differ significantly from fed controls (Fig. 2F). This inhibition of GSK3β by phosphorylation is expected to prevent desmin filament dissociation and myofibril breakdown during fasting (Kirk et al., 2014; Aweida et al., 2018). Accordingly, the major atrophy-related genes MuRF1 and MAFbx/Atrogin1, were strongly induced in skeletal muscle during fasting but not in the heart (Fig. 2G). Therefore, cardiac resistance to fasting-induced atrophy is associated with preservation of plakoglobin-containing membrane complexes and anabolic signaling.

### Plakoglobin supports force generation at increased sarcomere length

The localization of plakoglobin at membrane-cytoskeleton interfaces suggested that it may influence muscle mechanics. To test this idea, we measured contractile properties in skinned skeletal muscle fibers following plakoglobin knockdown. Plakoglobin was downregulated in mice by systemic intravenous injection of adeno-associated virus serotype 9 encoding a specific shRNA against the mouse plakoglobin gene *JUP* (AAV9.shJUP), while control mice received AAV9 encoding scrambled shRNA (AAV9.shLacz) (Fig. 3A). These constructs also encoded GFP to enable identification of infected muscle fibers. Robust GFP expression and reduced plakoglobin expression in skeletal muscle confirmed viral delivery and knockdown (Fig. 3B).

**Figure 3.**
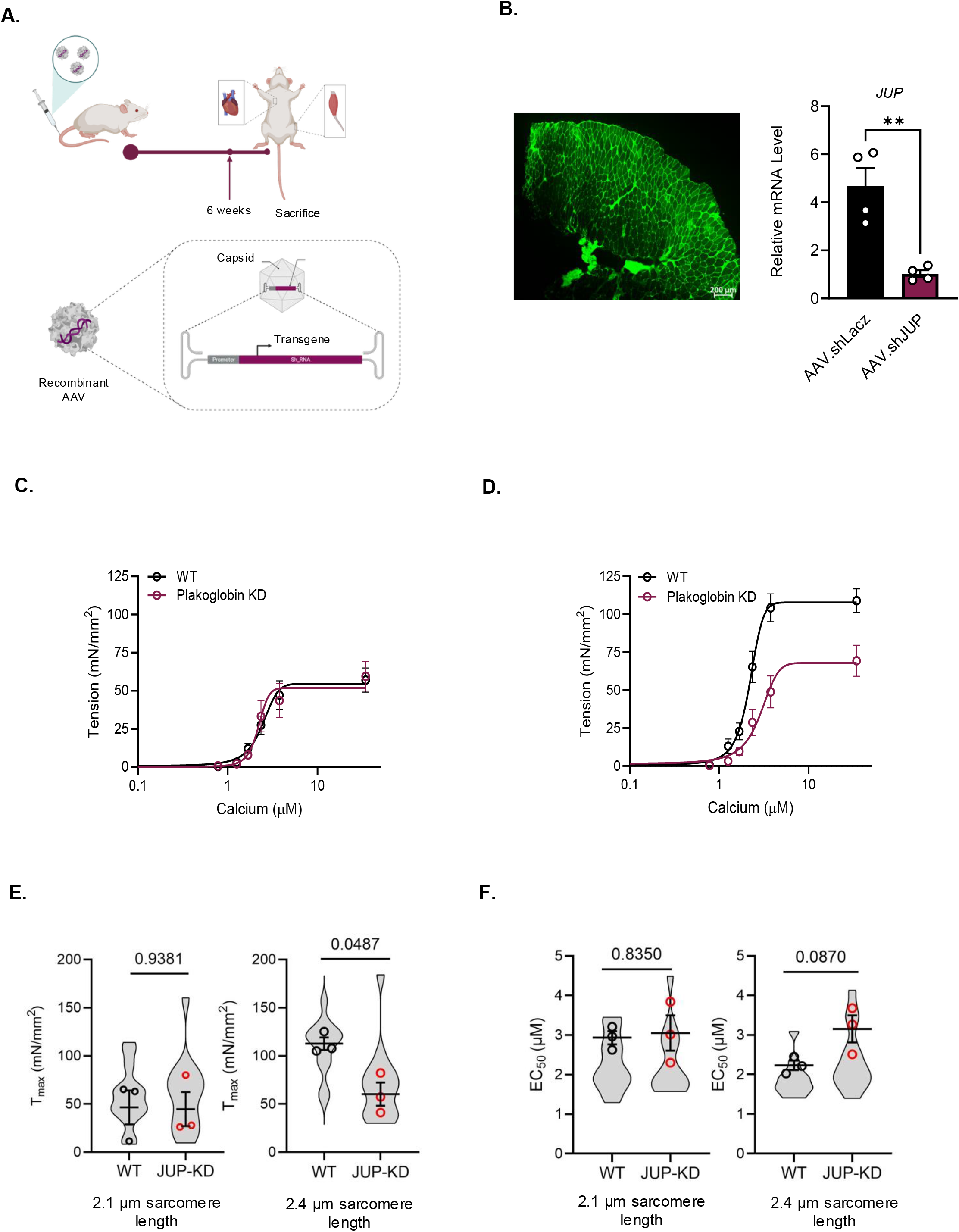
Plakoglobin is required for length-dependent activation of muscle fibers. **(A)** Schematic representation of systemic AAV administration in mice. Six weeks after AAV injection animals were sacrificed and muscles were dissected. **(B)** Efficient AAV delivery into mouse skeletal muscle. Left: TA muscle sections from AAV injected mice were stained with antibodies against eGFP (bar, 200 μm). Right: RT-PCR of mRNA preparations from AAV-infected muscles using specific primers for plakoglobin (*JUP)*. n = 4 mice per condition. Data are mean ± SEM. **, p<0.01 *vs*. shLacz by two-tailed unpaired Student *t* test. (C) Force–calcium curves of skinned skeletal muscle fibers from WT and plakoglobin knockdown muscle fibers at 2.1 µm sarcomere length. n = 3 mice per condition. (D) Force–calcium curves of skinned skeletal muscle fibers from WT and plakoglobin KD muscle fibers at 2.4 µm sarcomere length. n = 3 mice per condition. (E-F). Maximum calcium-activated tension (T_max_; E) and calcium sensitivity (EC_50_; F) of skeletal muscle fibers were analyzed. For WT, n=14 fibers at 2.1 μm and n=13 fibers at 2.4 μm; for plakoglobin knockdown (JUP-KD), n= 15 fibers at each sarcomere length. Fibers were obtained from n = 3 mice per group. Violin plots show individual fiber measurements; open circles represent the mean for each mouse, and horizontal lines and error bars represent mean ± SEM. WT and JUP-KD fibers were compared separately at each sarcomere length using two-sided nested *t* tests implemented as linear mixed-effects models, with genotype as a fixed effect and mouse as a random effect to account for multiple fibers measured within each mouse. Exact p values are shown.

Single skinned fibers were isolated from infected muscles, attached via micropins to a force transducer (Aurora Scientific) and length controller (Thor Labs), and exposed to increasing calcium concentrations while tension generation was measured at different sarcomere lengths of 2.1 and 2.4 μm. At 2.1 μm, the tension-calcium relationship was nearly identical between the control and the plakoglobin-deficient fibers (Fig. 3C). However, when the same experiment was performed at a long sarcomere length (2.4 μm), the shLacz expressing control fibers exhibited an increase in tension development, compared to the short sarcomere length, that was completely absent in plakoglobin-deficient fibers (Fig. 3D). Quantification of maximal tension confirmed these observations. At 2.1 μm, maximal calcium-activated tension (T_max_) did not differ significantly between control and plakoglobin-deficient fibers (Fig. 3E). However, when the muscle was stretched to 2.4 μm, fibers lacking plakoglobin generated significantly less T_max_ compared with control fibers (Fig. 3E). Thus, plakoglobin depletion selectively impaired force generation under conditions in which sarcomere length and mechanical load are increased.

To determine whether plakoglobin affects the calcium responsiveness of the contractile apparatus, we analyzed the calcium sensitivity of contraction (calcium concentration that generated half-maximal tension, EC_50_). Calcium sensitivity did not differ significantly between control and plakoglobin-deficient fibers at either sarcomere length (Fig. 3F). These findings indicate that plakoglobin is required for the enhanced contractile activation of skeletal muscle upon stretch.

### Cardiac muscle resists AAV-shJUP-mediated loss of plakoglobin

In light of these findings and our previous ones (Eid Mutlak et al., 2020) demonstrating that plakoglobin-containing protein assemblies contribute to maintenance of muscle integrity, we investigated whether downregulation of plakoglobin alone would be sufficient to trigger cardiac muscle atrophy *in vivo*. In TA muscle, AAV9.shJUP reduced plakoglobin mRNA and protein (Fig. 4A and B), decreased the abundance of β-dystroglycan and insulin receptor-AKT signaling (Fig. 4A), and increased MuRF1 expression (Fig. 4B). These effects reproduce the established response to plakoglobin loss in skeletal muscle, where *JUP* down-regulation causes atrophy (Cohen et al., 2014; Eid Mutlak et al., 2020) . In contrast, in the heart, plakoglobin abundance, insulin signaling, and MuRF1 expression remained similar to shLacz controls (Fig. 4B and C), despite shJUP viral delivery, as indicated by GFP expression (Fig. 4C). Thus, the heart resists AAV9.shJUP-mediated plakoglobin depletion, consistent with its ability to preserve plakoglobin complexes during fasting.

**Figure 4.**
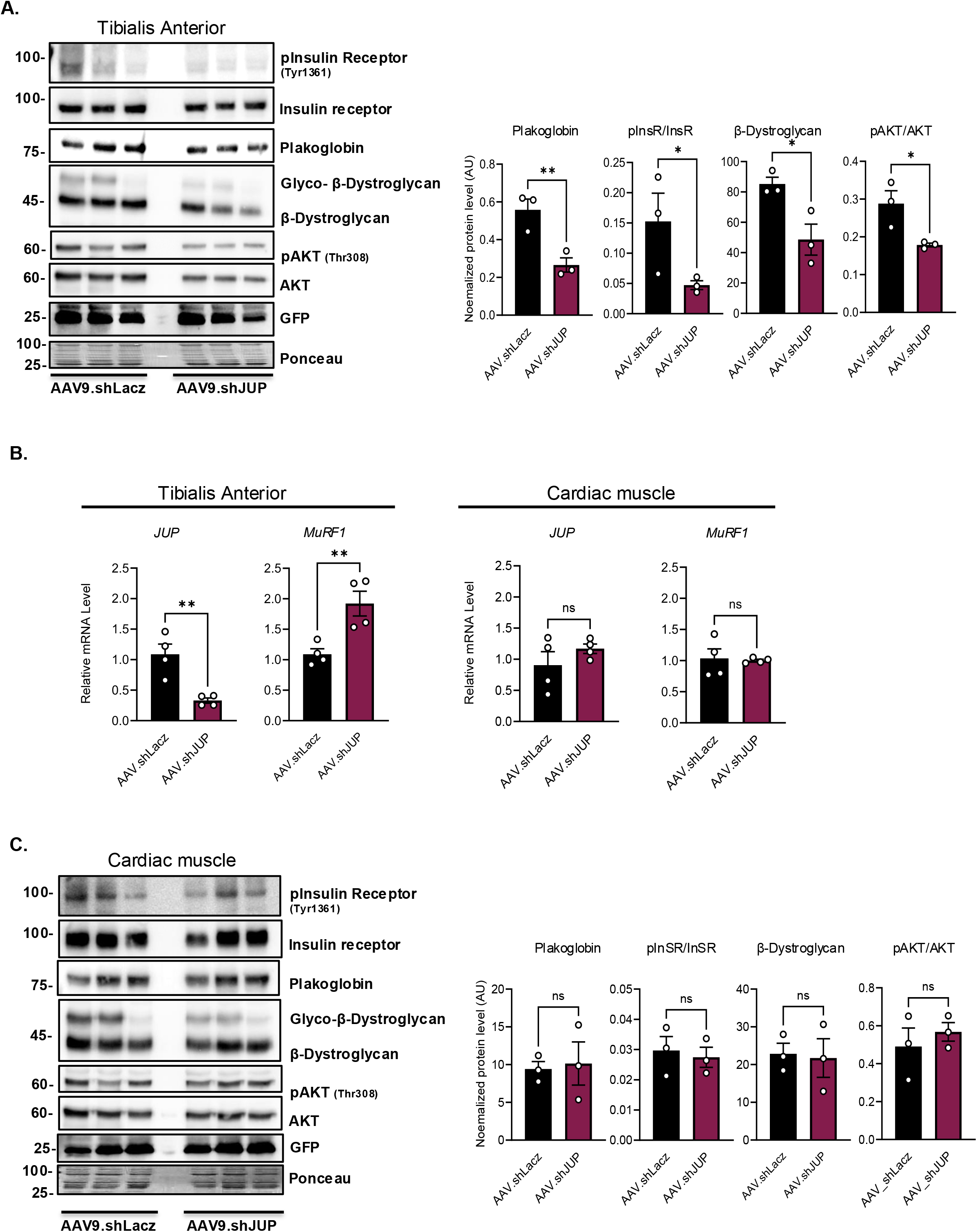
Cardiac muscle resists AAV9.shJUP-mediated depletion of plakoglobin. (A) Downregulation of plakoglobin reduces DGC stability and insulin signaling in skeletal muscle. Left: soluble fractions from AAV infected TA muscles were analyzed by immunoblotting. Right: densitometric measurements of the shown blots. n = 3 mice per condition. Data are mean ± SEM. *, p<0.05 and **, p<0.01 *vs*. shLacz by one-tailed unpaired Student *t* test. (B) Downregulation of plakoglobin in normal TA muscle activates MuRF1 expression, but not in the heart. RT-PCR of mRNA preparations from AAV-infected muscles using specific primers for plakoglobin (*JUP)* and MuRF1. n = 4 mice per condition. Data are mean ± SEM. ns, not statistically significant. **, p<0.01 *vs*. shLacz by two-tailed unpaired Student *t* test. (C) Left: soluble fractions from AAV infected cardiac muscles were analyzed by immunoblotting. Right: densitometric measurements of the shown blots. n = 3 mice per condition. Data are mean ± SEM. ns, not statistically significant *vs*. shLacz by two-tailed unpaired Student *t* test.

### MEK-ERK signaling promotes anabolic signaling and cardiac protection during fasting

The preservation of plakoglobin-associated membrane complexes and insulin signaling in cardiac muscle during fasting, when circulating insulin is low, suggested that additional cardioprotective mechanisms help maintain anabolic signaling in the heart. To identify pathways that may sustain cardiac signaling during fasting, we fractionated cardiac membrane-cytoskeleton extracts by gel filtration chromatography using a Superdex 200 10/300 column and analyzed fractions containing insulin receptor by mass spectrometry. Two fractions (#17 and #33) contained insulin receptor together with β-dystroglycan and plakoglobin (Fig. 5A). Network analysis of the proteins identified in these fractions revealed functional clusters of contractile proteins (red), DGC components (including dystrophin, dystroglycan and the core DGC component sarcoglycan (Wan et al., 2025)) and associated membrane structural proteins (blue), and proteins regulating MAPK signaling (green) (Fig. 5B). Among the identified components that regulate MAPK signaling were Tyrosine phosphatase PTPN11 (SHP2)(Scheiter et al., 2024), receptor for activated protein C kinase 1 (RACK1), a scaffold protein that organizes signaling complexes involving Src family kinases and MAPK signaling modules (Adams et al., 2011; McCahill et al., 2002), TRIP10, a Cdc42-interacting protein involved in membrane and cytoskeletal signaling (Hu et al., 2009), and G3BP1, a Ras-GTPase-associated protein connected with receptor signaling complexes (Yang et al., 2024) (Fig. 5B and Table 1). Their presence in insulin receptor-plakoglobin enriched fractions suggests that activation of Ras-MAPK pathway may contribute to maintenance of anabolic signaling in the heart. Consistently, immunoblot analysis of cardiac muscle homogenates from fed and fasted mice showed that phosphorylation of ERK1/2 increased during fasting (Fig. 5C).

**Figure 5.**
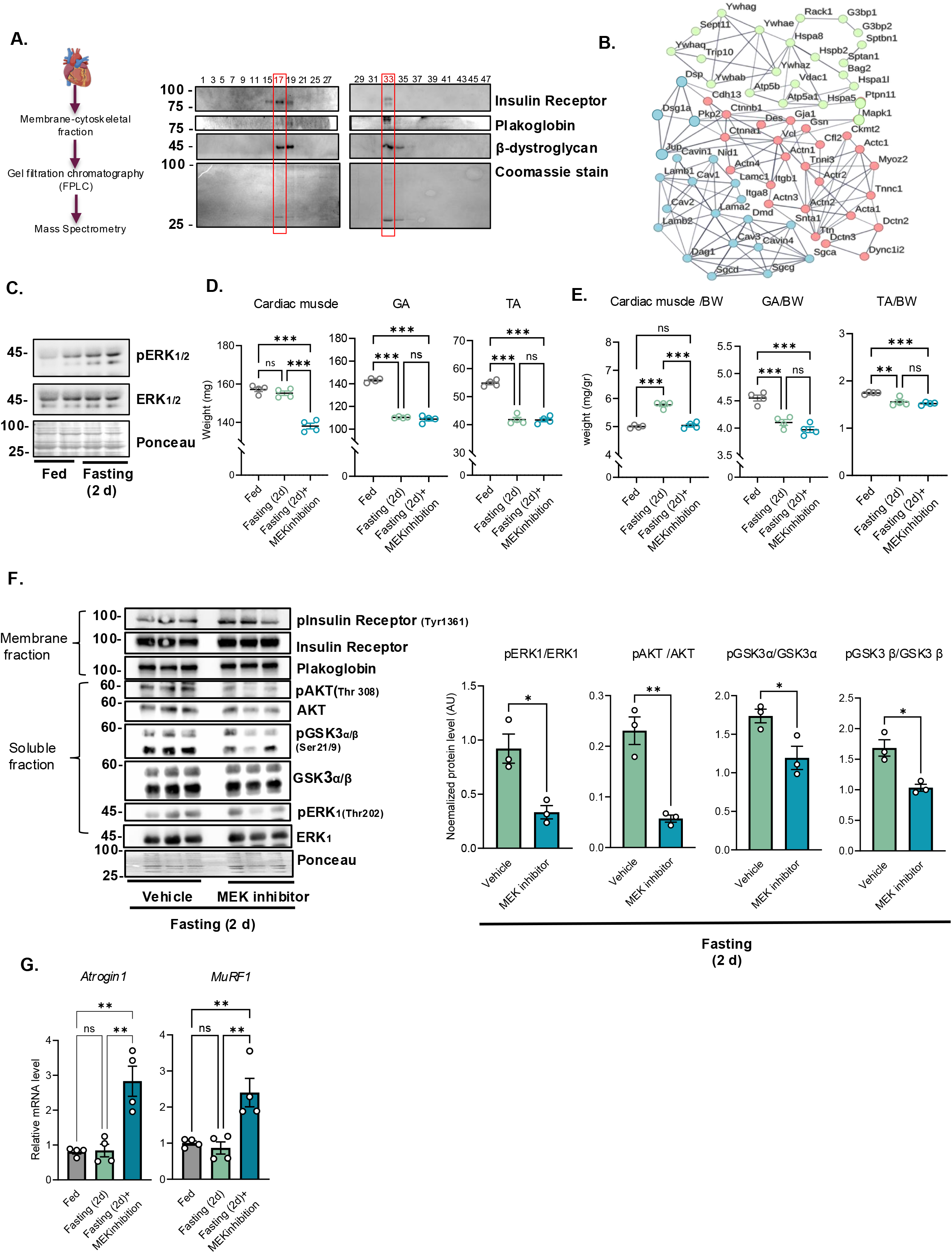
ERK signaling is required for cardiac muscle protection during fasting. (A) Left: Schematic of the FPLC experiment. A membrane–cytoskeletal extract from mouse cardiac muscle was fractionated by gel filtration chromatography using FPLC. Right: Analysis of eluted fractions by SDS–PAGE and immunoblotting using insulin receptor, plakoglobin and β-dystroglycan antibodies. Fractions containing all three proteins (#17 and #33) were analyzed by mass spectrometry. (B) Interaction network of components identified in proteomic analysis using the STRING database (version 12.0). Active interactions are based on published data, using interaction score of high confidence (0.7). (C) ERK signaling is enhanced in cardiac muscle during fasting. Immunoblot analysis of cardiac muscle homogenates from fed and fasted mice. n = 2 mice per condition. (D-E) During fasting, pharmacological MEK inhibition causes a reduction in cardiac muscle mass. (D) Mean muscle weights (mg) from fed and fasted (2 d) mice, and fasted mice injected with MEK inhibitor are depicted in graphs. (E) TA, GA and cardiac muscle weights are shown as ratios to BW (mg/gr). n = 4 mice per condition. Data are mean ± SEM. ns, not statistically significant, **, p<0.01 and ***, p<0.001 by one way ANOVA followed by Tukey’s post hoc multiple-comparisons test. (F) MEK inhibition during fasting reduces insulin signaling in the cardiac muscles. Immunoblot analysis of cardiac muscle membrane and soluble fractions from fasted mice injected with vehicle or MEK inhibitor. Right: densitometric measurements of the presented blots. n = 3 mice per condition. Data are mean ± SEM. *, p<0.05 and **, p< 0.01 *vs*. vehicle by two-tailed unpaired Student *t* test. (G) MEK inhibition during fasting induces atrogene expression in cardiac muscle. RT-PCR of mRNA preparations from cardiac muscles using specific primers for MAFbx/Atrogin1 and MuRF1. n = 4 mice per condition. Data are mean ± SEM. ns, not statistically significant. **, p<0.01 by one way ANOVA followed by Tukey’s post hoc multiple-comparisons test.

**Table 1.** Proteins that were co-purified with insulin receptor from cardiac muscle.

| **Entry** | **Protein names** | **Gene Names** | **No. unique peptides in fraction 17** | **No. unique peptides in fraction 33** |
| --- | --- | --- | --- | --- |
| P63085 | Mitogen-activated protein kinase 1 | *MAPK1* | 2 |  |
| P97855 | Ras GTPase-activating protein-binding protein 1 | *G3BP1* | 4 | 3 |
| Q8CJ53 | Cdc42-interacting protein 4 | *TRIP10* | 8 | 4 |
| P68040 | RACK1 | *RACK1* | 13 | 7 |
| P97379 | Ras GTPase-activating protein-binding protein 2 | *G3BP2* |  | 1 |
| P35235 | Tyrosine-protein phosphatase non-receptor type 11 | *PTPN11* |  | 10 |
| Q9CQV8 | 14-3-3 protein beta/alpha | *YWHAB* | 4 |  |
| P68254 | 14-3-3 protein theta | *YWHAQ* | 4 |  |
| P62259 | 14-3-3 protein epsilon | *YWHA E* | 17 | 1 |
| P61982 | 14-3-3 protein gamma | *YWHAG* | 10 | 5 |
| P63101 | 14-3-3 protein zeta/delta | *YWHA Z* | 7 | 1 |
| P11531 | Dystrophin | *DMD* | 2 |  |
| P49817 | Caveolin-1 | *CAV1* | 5 |  |
| P51637 | Caveolin-3 | *CAV3* | 2 |  |
| Q9WVC3 | Caveolin-2 | *CAV2* | 2 |  |
| Q61234 | Alpha-1-syntrophin | *SNTA1* | 2 |  |
| P82350 | Alpha-sarcoglycan | *SGCA* | 2 |  |
| P82348 | Gamma-sarcoglycan | *SGCG* | 3 |  |
| P82347 | Delta-sarcoglycan | *SGCD* | 10 |  |
| Q62165 | Dystroglycan 1 | *DAG1* | 7 |  |
| P09055 | Integrin beta-1 | *ITGB1* | 5 |  |
| A2ARA8 | Integrin alpha-8 | *ITGA8* | 4 |  |
| Q9WTR5 | Cadherin-13 | *CDH13* | 3 |  |
| Q02257 | Junction plakoglobin | *JUP* | 10 | 15 |
| E9Q557 | Desmoplakin | *DSP* | 7 | 24 |
| P31001 | Desmin | *DES* |  | 3 |
| Q02248 | Catenin beta-1 | *CTNNB1* |  | 1 |

Native insulin receptor-containing protein complexes were isolated from membrane extracts from mouse hearts by gel filtration chromatography using FPLC. Proteins co-purified with insulin receptor were identified by mass spectrometry.

To determine whether MEK-ERK signaling protects cardiac muscle, mice were injected intraperitoneally with the MEK inhibitor U0126 during fasting, and the effects on skeletal and cardiac muscles were analyzed. As shown above (Fig. 2A), fasting caused a marked reduction in the mass of skeletal muscles, including the GA and TA, whereas cardiac muscle weight was relatively preserved (Fig. 5D). Systemic inhibition of MEK, however, reduced cardiac muscle mass compared with vehicle-treated fasted mice (Fig. 5D and E). Thus, MEK-ERK activity is required for preservation of cardiac muscle mass during fasting. MEK inhibition also reduced phosphorylation of AKT and GSK3α/β (Fig. 5F) and induced MuRF1 and MAFbx/Atrogin1 in the heart during fasting (Fig. 5G). Interestingly, plakoglobin and insulin receptor membrane content remained unchanged (Fig. 5F), indicating that MEK-ERK signaling is not required to maintain their abundance under these conditions. Therefore, MEK-ERK activity maintains anabolic signaling and suppresses the atrophy program in cardiac muscle during fasting, acting downstream of or in parallel with plakoglobin-containing membrane complexes.

## DISCUSSION

The present studies identify a role for plakoglobin in force generation at increased sarcomere length and a signaling response associated with cardiac resistance to fasting-induced atrophy. Plakoglobin downregulation reduced maximal force at the longer sarcomere length without producing a detectable difference in calcium sensitivity between genotypes. During fasting, the continuously contracting heart retained plakoglobin-containing membrane complexes and anabolic signaling and maintained its mass, whereas skeletal muscle lost these complexes and underwent atrophy. Moreover, cardiac plakoglobin resisted AAV-shJUP-mediated depletion despite viral delivery, and inhibition of MEK-ERK signaling induced the atrophy program and reduced cardiac mass. These findings extend plakoglobin function beyond its established role in insulin signaling and skeletal muscle size and identify preservation of plakoglobin-containing complexes as a feature of cardiac resistance to fasting-induced atrophy.

Super-resolution SIM imaging revealed plakoglobin puncta along the sarcolemma and in structures closely associated with desmin (Fig. 1). This spatial organization places plakoglobin at interfaces that connect the membrane with the contractile cytoskeleton. During fasting-induced skeletal muscle atrophy, plakoglobin was selectively lost from the sarcolemma, together with DGC-insulin receptor assemblies. Because plakoglobin associates with DGC components and remains spatially linked to desmin at the sarcolemma (Sciandra et al., 2023; Eid Mutlak et al., 2020), its loss from the sarcolemma may destabilize membrane signaling complexes and their connection to the cytoskeleton, thereby contributing to the structural and signaling changes that accompany muscle atrophy.

Plakoglobin downregulation did not alter maximal force at short sarcomere length but selectively abolished the normal increase in maximal tension observed upon stretch (Fig. 3). Thus, plakoglobin is required for maintaining contractile force under conditions of increased sarcomere length. Because the fibers were chemically skinned, the assay does not establish acute sensing by an intact sarcolemma. Instead, the phenotype may reflect persistent effects of plakoglobin loss on membrane-associated cytoskeletal organization and/or on the contractile apparatus. The molecular basis of length-dependent activation remains incompletely understood and is generally attributed to mechanical signals generated within the contractile apparatus and associated cytoskeletal networks. Plakoglobin associates with DGC components at costameres and remains spatially linked to the desmin intermediate filament network, which extends from the sarcolemma into the myofibrillar apparatus. This organization provides a potential route by which plakoglobin-containing complexes could influence the transmission of mechanical information between membrane-associated structures and sarcomeres, thereby modulating myofilament activation without directly participating in force generation. Importantly, in these experiments, stretch is measured directly as sarcomere length. Thus, the role of plakoglobin is not simply tethering membrane stretch to intracellular/sarcomere stretch, but a more complex interaction between the sarcolemma and individual sarcomere proteins that results in an enhanced response to the same amount of sarcomere stretch. Defining the molecular changes that persist after plakoglobin depletion will be important for establishing how plakoglobin supports force at increased sarcomere length.

This role of plakoglobin in force-dependent cellular responses is consistent with observations in other tissues. Structural analyses revealed that plakoglobin forms part of a lattice-like scaffold linking adhesion complexes to cytoskeletal networks within desmosomes (Al-Amoudi et al., 2011). In embryonic stem cells, plakoglobin links physical cues to developmental responses (Kohler et al., 2023), and in endothelial cells it transmits mechanical tension across VE-cadherin complexes (Uttekar et al., 2026). Together with our findings, these observations support a broader role for plakoglobin as a force-transmitting protein that links membrane-associated adhesion complexes to cytoskeletal force-transmission systems. The functional relevance of such membrane–cytoskeleton association is well established in muscle because disruption of the DGC destabilizes the sarcolemma and increases susceptibility to contraction-induced injury (Ibraghimov-Beskrovnaya et al., 1992; Wilson et al., 2022), whereas defective dystroglycan glycosylation weakens the linkage between muscle fibers and the extracellular matrix (Muntoni et al., 2011). We previously showed that plakoglobin associates with DGC components and promotes insulin receptor signaling in skeletal muscle (Eid Mutlak et al., 2020). The present findings extend this model and suggest that plakoglobin-containing complexes not only stabilize anabolic signaling assemblies but also participate in sensing and transmitting mechanical information. Consistently with this interpretation, plakoglobin is an essential component of desmosomal junctions that protect tissues exposed to chronic mechanical stress, including the heart and the epidermis, and mutations in plakoglobin cause inherited cardiomyopathies in humans.

The distinct responses of skeletal and cardiac muscle to fasting further link the preservation of plakoglobin-containing membrane complexes with maintenance of muscle mass. Skeletal muscle lost these complexes and insulin signaling, induced MuRF1 and MAFbx/Atrogin1, and underwent atrophy. In contrast, the heart retained plakoglobin-containing complexes and anabolic signaling, did not induce either atrogene, and maintained its mass. These coordinated responses are consistent with our previous demonstration that loss of plakoglobin-dependent insulin signaling causes skeletal muscle atrophy (Cohen et al., 2014; Eid Mutlak et al., 2020). Moreover, although GFP expression confirmed successful delivery and expression of the same AAV9.shJUP vector in both cardiac and skeletal muscle, plakoglobin was reduced only in skeletal muscle. This resistance to plakoglobin depletion suggests that the heart possesses mechanisms that maintain its abundance. Because the heart remains continuously active during fasting, the forces generated by contraction may contribute to the retention or stabilization of plakoglobin at the cardiomyocyte membrane, although the underlying mechanism remains to be determined.

Our proteomic analyses of insulin receptor-enriched fractions from cardiac muscle identified several proteins connected to MAPK signaling, including RACK1, a scaffold known to organize MAPK signaling complexes (Adams et al., 2011), and PTPN11 (SHP2), an activator of Ras-MAPK signaling downstream of receptor tyrosine kinases and integrins (Chen et al., 2024). ERK phosphorylation also increased in cardiac muscle during fasting (Fig. 5). Most importantly, pharmacological inhibition of MEK reduced AKT-GSK3 signaling, induced MuRF1 and MAFbx/Atrogin1, and decreased cardiac muscle mass. Thus, MEK-ERK activity is required to suppress the atrophy program and maintain cardiac muscle mass during fasting. Because MEK inhibition did not reduce plakoglobin membrane content, ERK activity is not required to preserve plakoglobin abundance at the sarcolemma under these conditions. Because cardiac plakoglobin resisted AAV-shJUP-mediated depletion, the necessity of plakoglobin for cardiac protection could not be tested directly. Our findings therefore identify preservation of plakoglobin-containing complexes and activation of MEK-ERK signaling as coordinated features of the cardiac response to fasting, with neither pathway sufficient on its own, while the functional relationship between them remains to be established.

ERK signaling can respond to mechanical and receptor-mediated inputs and regulate cardiac muscle growth and adaptation to stress (Bueno et al., 2000; Kehat and Molkentin, 2010). Reduced membrane tension promotes ERK activation through FGF receptor endocytosis during stem cell differentiation (De Belly et al., 2021), whereas mechanical stretch activates ERK signaling through epidermal growth factor receptor (EGFR)-dependent pathways and drives a mechanochemical feedback loop that coordinate tissue organization (Hino et al., 2020). In this study, ERK-associated proteins were co-purified with the insulin receptor and plakoglobin from cardiac muscle, and systemic MEK inhibition demonstrated that MEK-ERK activity contributes to maintenance of cardiac muscle mass and anabolic signaling during fasting. Because the heart remains continuously active during nutrient deprivation, contraction may contribute to the retention or stabilization of plakoglobin and to sustained ERK signaling. However, cell-type-specific perturbations and temporal measurements will be needed to determine whether cardiac contraction, plakoglobin retention, and ERK activation are causally linked.. The implications of this mechanism may also be relevant to disease. Mutations in the *JUP* gene cause arrhythmogenic right ventricular cardiomyopathy, which is generally attributed to defective mechanical coupling between cardiomyocytes (Li et al., 2011). Whether disruption of plakoglobin-dependent mechanotransduction or altered signaling through the pathways described here contribute to disease pathogenesis are intriguing questions for future research.

Collectively, these studies identify a role for plakoglobin in force generation at increased sarcomere length and distinguish the cardiac response to fasting from that of skeletal muscle. In the heart during fasting, preservation of plakoglobin-containing complexes coincides with sustained anabolic signaling, suppression of the atrophy program, and maintenance of muscle mass. Systemic MEK inhibition compromises this protection. Additionally, mechanical activity may help preserve plakoglobin at the sarcolemma and sustain ERK-AKT signaling (Singh et al., 2026). We propose that plakoglobin-containing membrane-cytoskeleton complexes integrate mechanical activity with signaling pathways that preserve striated muscle during catabolic stress.

## MATERIALS AND METHODS

### Animal work

All animal experiments were approved by the Technion Institutional Animal Care and Use Committee and conducted in accordance with the ethics guidelines of the Israel Council on Animal Experiments. Male Hsd:ICR mice (∼ 30 g; Envigo) were used in all experiments and housed in a temperature- and humidity-controlled facility under a 12-hour light/dark cycle. Routine care and monitoring were provided by trained personnel in the institutional animal facility. Animals were randomly assigned to experimental groups. In fasting experiments, food was removed from cages for 48 hours while water was provided ad libitum.

### AAV administration

AAV vectors encoding shRNA targeting scrambled shLacz or the 3’UTR of mouse plakoglobin (*JUP*; gene ID 16480) under U6 promoter (experimentally validated shRNA sequence by manufacturer to knock down plakoglobin in cultured cells) were purchased from Vector BioLabs (Malvern, PA, USA) (Cat shAAV-262554) and were of Serotype 9, a serotype with a proven tropism for skeletal muscle cells (Katwal et al., 2013). Viral stock titers were provided by the manufacturer. Viral particles (1× 10^12^ gc) were administered to mice via tail vein injection (150 μl).

### MEK inhibitor administration

To inhibit MEK activity, mice were treated with U0126 (Sigma, U120) at 10 mg/kg body weight or vehicle control (5% DMSO in saline) by intraperitoneal injection. Injections were administered twice, at the time of food deprivation and 24 hours later. No adverse effects were observed at the administered dose.

### Immunofluorescence staining and super resolution microscopy

TA muscle longitudinal sections from fed and fasted mice were embedded in paraffin and then gradually rehydrated in ethanol/PBS. For immunofluorescence, 1:50 dilution of anti-Plakoglobin (Acris, Cat# AP33204SU-N) and anti-Desmin antibody (developed by D.A. Fischman (Cornell University Medical College, New York, NY) was obtained from Developmental Studies Hybridoma Bank, created by the NICHD (University of Iowa, Iowa City, IA) (Cat# D3), and 1:1,000 dilution of Alexa Fluor 568 (Thermo Fisher Scientific, Cat # A-11075) and 647 (abcam, Cat# ab150115)–conjugated secondary antibody were used.

Muscle sections were imaged using the Elyra7 lattice SIM super resolution microscope (Zeiss) with a Plan-Apochromat 63x Oil objective (NA 1.46), 561nm and 488 nm lasers and a pco.edge sCMOS camera. Images were acquired using identical acquisition settings and 16-bit 2D image datasets were collected. Super-resolution reconstruction was performed using SIM^2 reconstruction algorithm and ZEN 3.3 black software (Zeiss). Image analysis was performed using the Imaris software (Bitplane, ver 9.31). Identical parameter settings were applied across all experimental groups and a consistent threshold was applied across all datasets. At least 2 random areas per mouse muscle section were analyzed and averaged (n=3 mice per group). Membrane regions of interest (ROIs) were defined by manual surface segmentation followed by channel masking. Stained protein puncta were segmented using the ‘Spots’ module. The spots were set to 100 nm in diameter that is below the light diffraction limit but above the SIM^2 super resolution imaging limit (60 nm). Spot counts for each image were obtained through the statistics tab. Colocalization measurements were performed using “Coloc Spots” extension, with a distance threshold between close spots was set to 100 nm from each other. All quantified data were exported directly from Imaris without manual post-processing.

For Fig. 3B, frozen TA muscle cross-sections were cut at 10 µm, fixed in 4% PFA for 20 min at room temperature, and washed three times in PBST (PBS, 0.1% Triton X-100). Sections were incubated in blocking solution (0.2% BSA, 5% goat serum in PBST) for 1 h at room temperature, stained with anti-GFP antibody (ab290, Abcam, 1:500) overnight at 4°C, and incubated with Alexa Fluor 488-conjugated secondary antibody (Abcam, #Cat ab150077, 1:1000) for 1 h at room temperature. Images were collected using an LSM 710 confocal laser scanning microscope (Zeiss) with a Plan-Apochromat 20×/0.8 M27 objective and a 488 nm laser. Fluorescence was detected using a photomultiplier tube (PMT). Images were acquired using identical acquisition settings, and 8-bit 2D image datasets were collected.

### Muscle homogenization

Muscles were homogenized in ice-cold lysis buffer (20 mM Tris, pH 7.6, 5 mM EGTA, 100 mM KCl, 1% Triton X-100, 1 mM PMSF, 10 mM sodium pyrophosphate, 3 mM benzamidine, 10 μg/ml leupeptin, 10 μg/ml aprotinin, 50 mM NaF, and 2 mM sodium orthovanadate) using an Omni Tissue Homogenizer and incubated at 4°C for 1hour. After centrifugation at 6,000 × g for 20 min at 4°C, the supernatant was stored at -80°C as the soluble fraction. All steps were performed on ice or at 4°C to preserve protein integrity.

### Muscle fractionation

To obtain membrane extracts, muscles were homogenized in ice-cold buffer C (20 mM Tris, pH 7.6, 100 mM KCl, 5 mM EDTA, 1 mM DTT, 1 mM PMSF, 3mM benzamidine,10μg/ml leupeptin, 50 mM NaF, and 1 mM sodium orthovanadate) using short mechanical bursts on ice. Homogenates were centrifuged at 2,900 × g for 20 min at 4°C to remove cell debris and nuclei. The supernatant (containing soluble cytosolic and membrane components) was collected and subjected to ultracentrifugation at 180,000 × g for 90 min at 4°C to isolate membranes. The obtained supernatant was stored as the cytosolic fraction. The pellet (containing membranes and associated proteins), was resuspended in ice-cold Buffer M (20 mM Tris-HCl, pH 7.6, 5 mM EDTA, 100 mM KCl, 1 mM DTT, 0.25% sodium deoxycholate, 1% NP-40, 1 mM sodium orthovanadate, 10 μg/ml leupeptin, 3 mM benzamidine, 1 mM PMSF, and 50 mM NaF), followed by gentle rotation for 20 min at 4 °C. After a second ultracentrifugation at 100,000 x g for 30 min, the supernatant was collected as the membrane-enriched protein fraction. All fractions were stored at -80 °C until further analysis. Protein concentration was determined using DC Assay Protein Assay Kit (Bio-Rad Laboratories, cat. #5000116) according to the manufacturer’s instructions.

### Co-Immunoprecipitation

Membrane extracts (100-200 μg) were incubated overnight at 4 °C with 1 μg of primary antibody per 100 μg of sample, or with an equivalent amount of control non-specific IgG, with rotation. Then, 40 μl of 50% pre-washed Protein A/G agarose slurry (Santa Cruz Biotechnology, sc-2003) was added and incubated for 3-4 hours at 4 °C with rotation. Beads were pelleted by centrifugation at 2,500 rpm for 5 min at 4 °C and washed twice with each of the following buffers: high-salt buffer (50 mM Tris-HCl, pH 8.0, 500 mM NaCl, 0.1% SDS, 0.1% Triton X-100, 5 mM EDTA), medium-salt buffer (same as above with 150 mM NaCl), and low-salt buffer (50 mM Tris-HCl, pH 8.0, 0.1% Triton X-100, 5 mM EDTA), with centrifugation at 2,500 rpm for 5 min at 4 °C between washes. After the final wash, beads were resuspended in SDS sample loading buffer, heated at 70 °C for 10 min, and analyzed by SDS-PAGE and immunoblotting.

### Western blotting and antibodies

Equal amounts of protein were resolved by SDS-PAGE (10 μg for membrane fractions and 25 μg for whole-cell lysates) and transferred to PVDF membranes (2 hours, 200mA, 4°C). Membranes were blocked for 1 hour in 5% BSA/TBS-T (20 mM Tris-HCl, pH 7.6, 150 mM NaCl, 0.1% Tween-20) and incubated with primary antibodies overnight at 4°C, washed in TBST, and incubated in secondary antibody for 1 hour at room temperature. Signals were detected using ECL reagent and the Fusion FX (Vilber) or Fusion solo (Vilber) imager systems. Protein band intensity was quantified by densitometric analysis of the immunoblots using Quantity One 1-D Analysis Software (Bio-Rad).

Plakoglobin antibodies were from Genetex (Cat# GTX15153, for immunoprecipitation (1:1000) and for immunoblotting (1:1000)), BD Transduction Laboratories (Cat# 610254, 1:1000). and anti-GFP (Cat# ab290, 1:1000) were from Abcam. Anti-β-dystroglycan antibody was developed by Morris, G.E. (RJAH Orthopedic hospital, Oswestry, UK) was obtained from the Developmental Studies Hybridoma Bank, created by the NICHD (University of Iowa, Iowa City, IA) (Cat# MANDAG2,7D11, 1:100). Anti-phospho-insulin receptor (Cat# 3023, 1:1000), anti-insulin receptor (Cat# 3025, 1:1000), anti-phospho-AKT (Cat#13038, 1:1000), anti-AKT (Cat# #4685, 1:1000) anti-Phospho-GSK-3 alpha/beta (cat# 8566, 1:1000), anti-GSK-3 alpha/beta (cat# 5676, 1:1000), anti-phospho-ERK (Cat# 9101, 1:1000) and anti-ERK (Cat# 4695, 1:1000) were from Cell Signaling Technologies. Syntrophin antibody (Cat# sc-166207, 1:1000), was from Santa Cruz Biotechnology. The secondary antibodies used in this study were - Peroxidase Goat Anti-Rabbit IgG (H+L) (Cat. #111-035-144, 1:10,000) and Peroxidase Goat Anti-Mouse IgG (H+L) (Cat. #115-035-003, 1:5,000) were from Jackson Immuno Research.

### Quantitative real-time PCR

Total RNA was extracted from muscles using TRIzol reagent (Cat. #T9424; Sigma-Aldrich) and served as a platform for cDNA synthesis using qScript cDNA Synthesis Kit (cat. #95047; QuantaBio). Real-time PCR was performed with primers to mouse genes (see Supplementary Table 1) using qPCRBIO SyGreen Blue Mix Hi-ROX (Cat. #PB20.16-05; PCR BIOSYSTEMS), according to manufacturer’s instructions. The gene GAPDH was used as a reference.

### Isolation of native insulin receptor complexes from cardiac muscle

Normal mouse cardiac muscle was homogenized in 19 volumes of ice-cold buffer A (20 mM Tris, pH 7.2, 100 mM KCl, 5 mM EGTA, 40 mM imidazole, 1 mM dithiothreitol (DTT), 1 mM phenylmethylsulfonyl fluoride (PMSF), 3 mM benzamidine, 10 μg/ml leupeptin, 50 mM NaF, 2.7 mM sodium orthovanadate, and 1% Triton X-100). After centrifugation at 10,000 × g for 20 min at 4 °C, the pellet (containing membranes, cytoskeleton, and myofibrils) was washed once in buffer A, and then re-suspended in 10 volumes (v/v) of buffer B (buffer A supplemented with 10 mM ATP) to dissociate myofibrillar myosin from actin thin filaments. After centrifugation at 16,000 × g for 10 min at 4 °C, the pellet (mostly containing membrane and cytoskeleton-bound proteins) was washed once in buffer B, re-suspended in 10 volumes (v/v) of buffer C (buffer A at pH 9), and incubated at 37 °C for 30 min with gentle agitation. After centrifugation at 16,000 × g for 10 min and 4 °C, the supernatant (containing purified membrane-bound and cytoskeletal proteins) was subjected to 30% ammonium sulfate precipitation, and precipitates were centrifuged at 20,000 × g for 30 min at 4 °C. The obtained pellet was kept on ice, and the supernatant was subjected to an additional 50% ammonium sulfate precipitation. After a final centrifugation at 20,000 × g for 30 min at 4 °C, precipitates from 30% and 50% ammonium sulfate precipitations were re- suspended in 600 µl of buffer A, and 85% of this suspension (containing membrane-cytoskeleton fraction) was loaded onto a Superdex 200 10/300 Gel Filtration column (GE Healthcare Life Sciences, UK) equilibrated with 20 mM Tris, pH 7.2. Proteins were eluted at a flow rate of 0.3 ml/min, and 300 μl fractions were collected. Selected fractions enriched with insulin receptor, plakoglobin and β-dystroglycan (Fig. 5A) were analyzed by mass spectrometry and western blotting.

### Mass spectrometry analysis

The mass spectrometry data were analyzed using Proteome Discoverer 2.4 (Thermo) using Sequest search engine, searching against the Mus musculus proteome from the Uniprot database with mass tolerance of 20 ppm for the precursor masses and 0.02 Da for the fragment ions. Oxidation on methionine, and protein N-terminus acetylation were accepted as variable modifications and carbamidomethyl on cysteine was accepted as static modifications. Minimal peptide length was set to six amino acids and a maximum of two miscleavages was allowed. The data was quantified by label free analysis using the same software. Peptide-level false discovery rates (FDRs) were filtered to 1% using the target-decoy strategy.

### Single Skinned Muscle Fibers

Skinned muscle fibers were prepared by homogenizing GA muscles with three 1 second pulses at 7000 rpm in an isolating solution containing Triton X-100, protease and phosphatase inhibitors. Following a 20-min incubation, the sample was centrifuged at 120 x g for 2 min at 4 °C and repeated twice after resuspension in isolating solution. Using an Aurora Scientific force transducer and Piezo length controller, a single skeletal muscle fiber was attached to two pins via UV glue (NOA68/NOA61, Thor Labs). The sarcomere length was measured to 2.1 µm and the myocyte was put into a relaxing solution with protease and phosphatase inhibitors. Calcium activated force measurements were taken at 2.1µm by moving the myocyte from the relaxing solution into baths containing different concentrations of calcium activating solution (100%, 90%, 85%, 80%, 75%, 65%). After all data points were collected, the cell was stretched to 2.4 µm and the experiment was repeated. The force was normalized to the myocyte cross-sectional area, and the data was fitted to a Hill Curve. All tests were completed at room temperature.

### Statistical Analysis

Statistical analyses and graph preparation were performed using GraphPad Prism 10 (GraphPad Software). Data are presented as mean ± SEM unless otherwise indicated. The experimental unit and sample size (n) for each analysis are indicated in figure legends. For animal experiments, n represents individual mice. When multiple measurements were obtained from the same mouse, the individual measurements were not treated as independent biological replicates. Comparisons between two independent groups were performed using two-tailed unpaired Student’s *t* tests. A one-tailed unpaired Student’s *t* test was used only in Figs. 2C and 4A because the directional hypothesis, that loss of plakoglobin reduces plakoglobin-associated membrane proteins and insulin signaling in skeletal muscle was established a priori based on our previous studies. Comparisons among three or more groups were performed using one-way ANOVA followed by Tukey’s multiple-comparisons test, as indicated in the figure legends. For the skinned-fiber experiments in Fig. 3E–F, WT and JUP-KD groups were compared at each sarcomere length using two-sided nested *t* tests implemented as linear mixed-effects models, with genotype as a fixed effect and mouse as a random effect to account for replicate fiber measurements within individual mice.

Statistical tests and p values are reported in the figure legends; p < 0.05 was considered statistically significant.

### Data availability

All datasets generated and analysed during this study, along with the materials and experimental protocols supporting the findings, are available from the corresponding author upon reasonable request. The mass spectrometry proteomics data generated in this study have been deposited in the ProteomeXchange Consortium via the PRIDE partner repository under the accession number PXD082518.

## ACKNOWLEDGMENTS

This study was funded by the European Union under 101080229-2 to S. Shemer. It was part of the DREAMS project. The funders played no role in study design, data collection, analysis and interpretation of data, or the writing of this manuscript. Views and opinions expressed are however those of the author only and do not necessarily reflect those of the European Union (EU) or European Research Executive Agency (REA). Neither the EU nor REA can be held responsible for them. Additional funds have been received from the Bruce and Ruth Rappaport Cancer Research Center and Hellen and Norman Asher fund to S. Shemer, to support the proteomics analysis at the Smoler Proteomics Center, Technion.

We thank Dr. Izhak Kehat for the constructive feedback and Dr. Nitsan Dahan for the kind assistance with SIM imaging of muscle sections. We also thank Dr. Anqi Chen and Dr. David A. Weitz for their contributions and helpful discussions during the early development of this project. The illustrations in Figs. 1A and 3A were created with BioRender.com.

## Author contributions

S.C, N.J. and S.M.D.: data curation, formal analysis, investigation, validation, project administration, paper writing. J.A.K. data curation, formal analysis, investigation, validation, manuscript writing and editing. S.S: validation, conceptualization, funding acquisition, project administration, supervision, manuscript writing and editing.

## SUPPLEMENTARY MATERIAL

**Supplementary Table 1.**
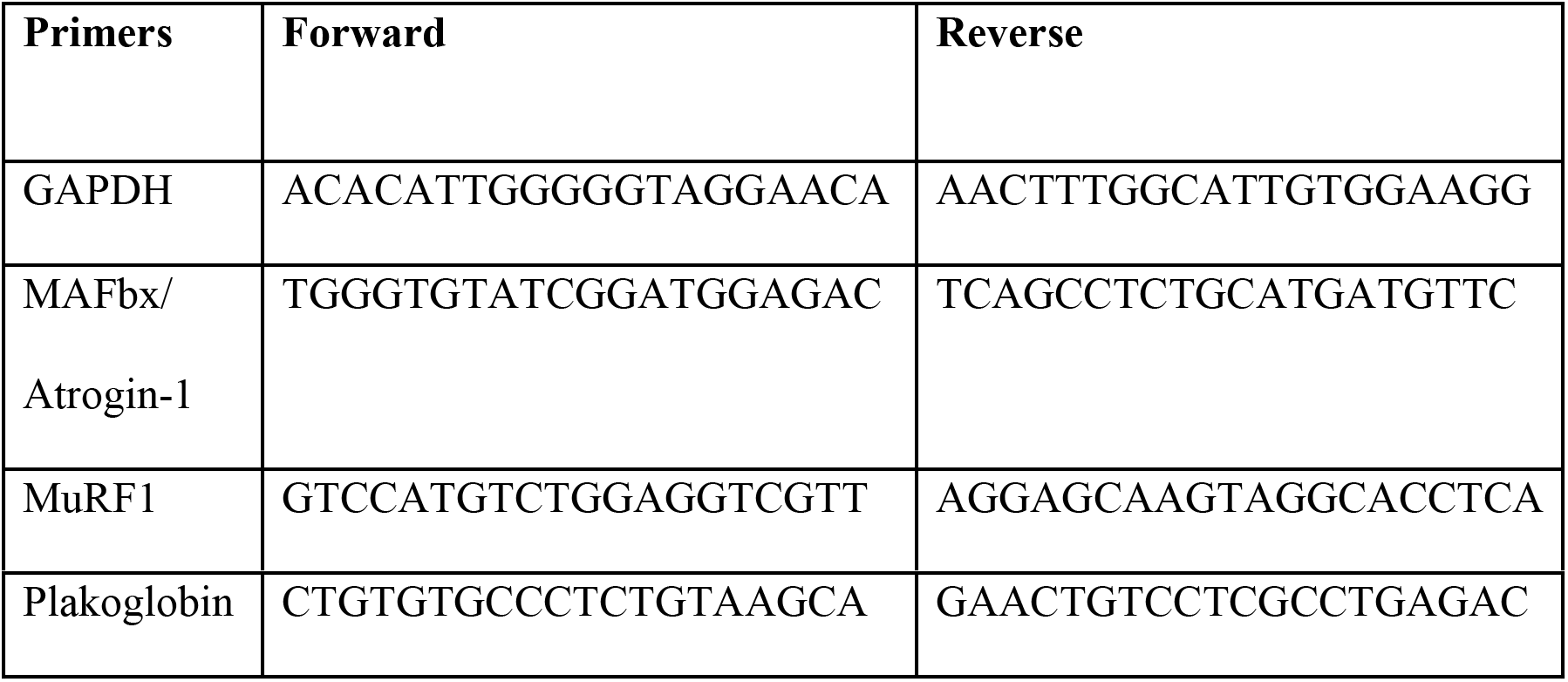
Quantitative PCR primers used in the present study.

